# Tetraeffective causes, mortacauses, and vitacauses of mortality and survivorship

**DOI:** 10.1101/039438

**Authors:** Michael Epelbaum

## Abstract

Every tetraeffective cause of mortality and survivorship negatively and positively affects mortality and negatively and positively affects survivorship. There is previous evidence of tetraeffective causes of mortality and survivorship, and strong rationales suggest that every cause of mortality and survivorship is tetraeffective. Here I elucidate and explain that every tetraeffective cause of mortality and survivorship combines corresponding at least one cause-specific mortacause and at least one cause-specific vitacause; “mortacause” refers here to a cause-specific component that positively affects mortality and negatively affects survivorship, and “vitacause” refers to a cause-specific component that positively affects survivorship and negatively affects mortality. I show tetraeffective causes, mortacauses, and vitacauses in results of multivariable regression analyses of effects of age, lifespan, contemporary aggregate size, lifespan aggregate size, and historical time humans’ and medflies’ mortality and survivorship. In these analyses I specify tetraeffective causes, mortacauses, and vitacauses with *sign(β_1_)* = -*sign(β_2_)*, where respective corresponding *β_1_* and *β_2_* denote respective first and second variable-specific regression coefficients. Thus tetraeffective causes, mortacauses, and vitacauses of mortality and survivorship are hereby defined, identified, named, recognized, elucidated, conceptualized, specified, explained, and demonstrated.

---

Wildfires are tetraeffective causes that negatively and positively affect mortality and survivorship of plants and animals^1^. There is also ample evidence of iatrogenic effects on mortality and survivorship^2,3^, including, for example, iatrogenic effects of surgery^4^ and pharmacologic medication (e.g., antibiotics^5,6^). In its totality, previous research on social, economic, cultural, or political causes of humans’ mortality and survivorship shows that each of these kinds of causes is tetraeffective^7-21^. Additionally, Strehler-Mildvan correlations^22-29^, compensations^27,28,30,31^, and hysteresis or delays^32-34^ in effects of age on mortality and survivorship show that age is a tetraeffective cause of mortality and survivorship. These considerations show that there is evidence of tetraeffective causes of mortality and survivorship. However, until now, tetraeffective causes of mortality and survivorship have remained undefined, unidentified, unnamed, unrecognized, unclear, misconceived, unspecified, and unexplained.

Here I elucidate and explain that every tetraeffective cause of mortality and survivorship combines corresponding at least one cause-specific mortacause and at least one cause-specific vitacause. Fig. 1 depicts the causal structure of a tetraeffective cause *X* that affects mortality *M* and survivorship *S* through the combined positive effects of an *X*-specific mortacause *Xm* on mortality *M*, negative effects of an *X*-specific mortacause *Xm* on survivorship *S*, negative effects of an *X*-specific vitacause *Xv* on mortality *M*, and positive effects of an *X*-specific vitacause *Xv* on survivorship *S* (e.g., illustrating that an increasing *Xm* leads to increasing *M* and decreasing *S* and illustrating that an increasing *Xv* leads to increasing *S* and decreasing *M*). Tetraeffective causes could have diverse causal structures; for example, while Fig. 1 presents a causal structure of a tetraeffective cause with one mortacause and one vitacause, Fig. 2 presents a causal structure of a tetraeffective cause with more than one mortacause and more than one vitacause. The causal structures that are depicted in Fig. 1 and Fig. 2 are consistent with the laws of identity, noncontradiction, and excluded middle^35-37^. In contrast, Fig. 3 depicts the causal structure of a cause *X* that directly negatively and positively affects mortality *M* and directly negatively and positively affects survivorship *S*. The causal structure that is depicted in Fig. 3 is inconsistent with the laws of identity, noncontradiction, and excluded middle; moreover, the causal structure that is depicted in Fig. 3 does not validly depict the causal structure of tetraeffective causes of mortality and survivorship.

**Figure 1:**
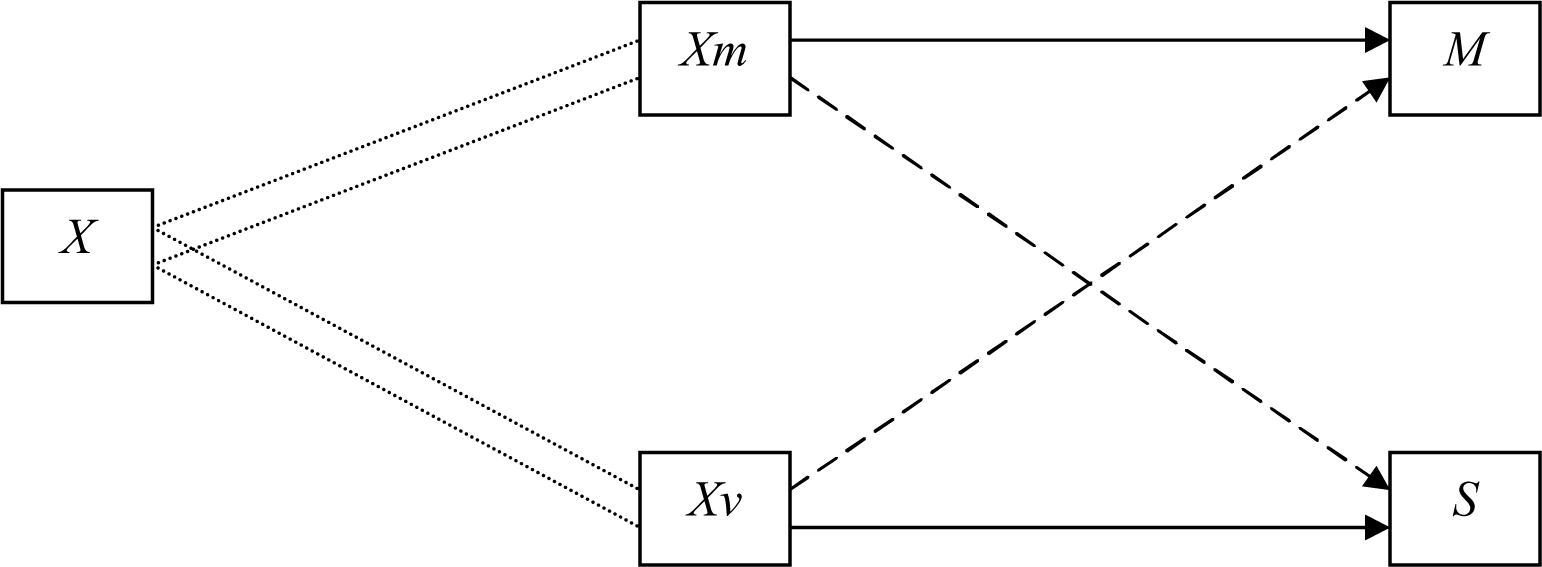
A causal structure of effects one mortacause and one vitacause on mortality and survivorship. *X* denotes a tetraeffective cause of mortality *M* and survivorship *S*, *Xm* denotes an *X*-specific mortacause, *Xv* denotes an *X*-specific vitacause, double dotted lines denote that *Xm* and *Xv* are *X*-specific, arrow → denotes positive effects, and arrow ----> denotes negative effects.

**Figure 2:**
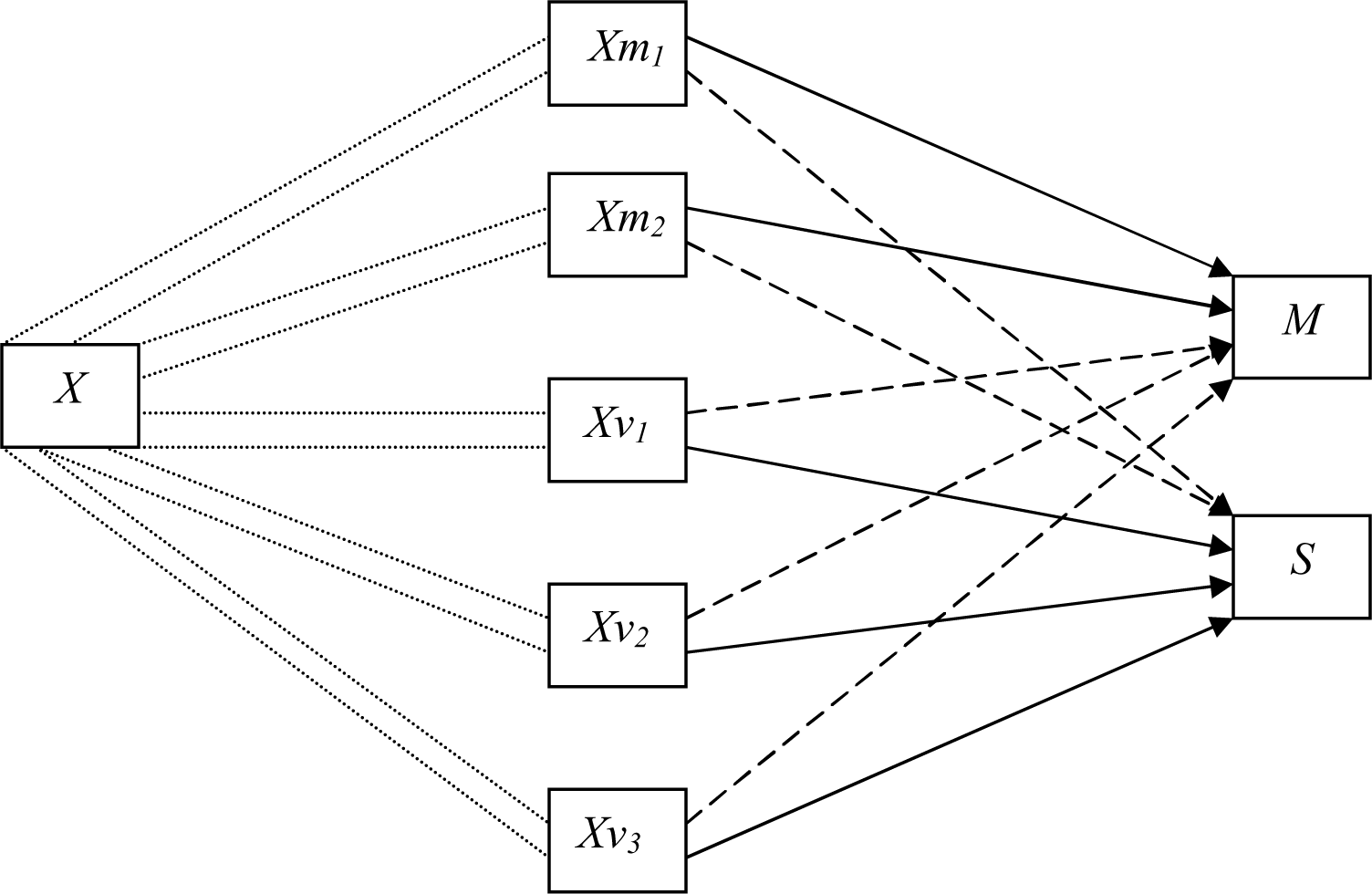
A causal structure of effects of two mortacauses and three vitacauses on mortality and survivorship. *X* denotes a tetraeffective cause of mortality *M* and survivorship *S*, *Xm* denotes an *X*-specific mortacause, *Xv* denotes an *X*-specific vitacause, double dotted lines denote that *Xm* and *Xv* are *X*-specific, arrow → denotes positive effects, and arrow ----> denotes negative effects.

**Figure 3:**
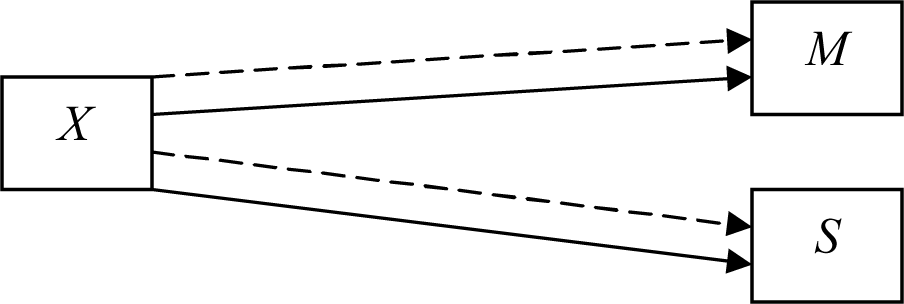
A causal structure of direct negative and positive effects on mortality and direct negative and positive effects on survivorship. *X* denotes a cause that directly negatively and positively affects mortality *M* and survivorship *S*, arrow → denotes positive effects, and arrow ----> denotes negative effects.

Mortality refers to cessation of existence, survivorship refers to continuation of existence, and that which exists or ceases to exist is an entity. An entity can be simple or complex, natural or artificial, living or non-living; a particle, droplet, cell, virus, insect, human, plant, rock, lake, mountain, planet, celestial system, universe, city, sculpture, bicycle, airplane, basketball team, nation state, language, book, or poem are some of the many examples of an entity. All previous entities existed and ceased to exist, the universe and all its present and future entities will cease to exist^38-41^, and continuations and cessations of existences are regulated^42-50^; therefore, mortality, survivorship, and their causes are regulated regulators of the existence of every entity. If every cause of mortality and survivorship is a tetraeffective cause of mortality and survivorship, then tetraeffective causes are intimately involved in the existence – and the continuation, regulation, and limitation of existence – of every entity. However, if diverse but not all causes of mortality and survivorship are tetraeffective causes of mortality and survivorship, then existence – and continuation, regulation, and limitation of existence – do not consistently apply to every entity. Similarly, if no causes of mortality and survivorship are tetraeffective, then existence – and continuation, regulation, and limitation of existence – do not consistently apply to every entity. Therefore, every cause of mortality and survivorship is tetraeffective.

The total number of causes of mortality and survivorship in an hypothetical system that involves only tetraeffective causes in the regulated regulations and limitations of existence of all entities is smaller than the total number of causes of mortality and survivorship in an hypothetical system that excludes tetraeffective causes of mortality and survivorship from the regulated regulations and limitations of existence of all entities. Therefore, a system that involves tetraeffective causes of mortality and survivorship in the regulated regulations and limitations of existence of all entities is more parsimonious than a corresponding system that excludes tetraeffective causes of mortality and survivorship. This parsimony provides an additional rationale for the universality of tetraeffective causes of mortality and survivorship. Furthermore, symbioses between corresponding at least one cause-specific mortacause and at least one cause-specific vitacause of every tetraeffective cause of mortality and survivorship are illustrated by the observation that any positive effect of age on mortality and any negative effect of age on survivorship require entities of ages greater than zero, and any entity of age greater than zero requires corresponding negative effects of age on mortality and positive effects of age on survivorship; these symbioses and requirements imply that effects of age on mortality and survivorship are tetraeffective, further implying that age is a tetraeffective cause of mortality and survivorship. Similar symbioses, requirements, and implications apply to every entity and every cause of mortality and survivorship. These considerations provide additional rationales for the universality of tetraeffective causes of mortality and survivorship.

Extensive and longstanding considerations of oppositions in religion and philosophy^35,36,51-63^, quantum theory^64,65^, structuralism^66-68^, biology^69-71^, and art^72-77^ imply that every mortacause is opposed by – and opposes – a corresponding at least one vitacause. These considerations also imply that every vitacause is opposed by – and opposes – a corresponding at least one mortacause. Additionally, these considerations imply that every tetraeffective cause opposes – and is opposed by – another tetraeffective cause. However, if every cause of mortality and survivorship is not tetraeffective (i.e., if tetraeffective causes of mortality and survivorship do not exist) then at least some causes of mortality are not opposed and at least some causes of survivorship are not opposed, such that these absences of oppositions violate the requisites of opposition. Moreover, if every cause of mortality and survivorship is not tetraeffective then at least some entities do not cease to exist, violating the law of cessation of existence of every entity. Therefore, and in consistency with ample previous scientific consideration of intrinsic and extrinsic causes of mortality and survivorship^25,27,31,78-91^, as well as in consistency with previous considerations of essential and coincidental properties^35,92^ – corresponding at least one mortacause and at least one vitacause must be intrinsic to every cause of mortality and survivorship and, therefore, every cause of mortality and survivorship is tetraeffective.

Components of tetraeffective causes of mortality and survivorship can be hidden (e.g., these components can be unknown, unobserved, ignored, or misconceived). However, the hiddenness of at least one mortacause of a tetraeffective cause of mortality and survivorship does not imply the following: (i) the at least one mortacause does not exist, and (ii) the cause is not tetraeffective. Similarly, the hiddenness of at least one vitacause of a tetraeffective cause of mortality and survivorship does not imply the following: (i) the at least one vitacause does not exist, and (ii) the cause is not tetraeffective. Therefore, it is invalid to conclude that a cause – e.g., every cause, any cause, a specific cause – of mortality and survivorship is not a tetraeffective cause. Furthermore, the continuation of an entity’s existence does not mean that respective causes of the cessation of this entity’s existence do not affect this entity’s existence; similarly, the cessation of an entity’s existence does not mean that respective causes of the continuation of this entity’s existence do not affect this entity’s cessation of existence. These considerations show that the universality of tetraeffective causes of mortality and survivorship is undeniable.

Diverse cultures, religions, philosophies, and scientific investigations consider effects of damage, frailty, disease, injury, waste, harm, poison, thanatos, destroyer of worlds, nuclear holocaust, global warming, poverty, injustice, or similar phenomena on mortality and survivorship^27,28,60-63,93-108^. Similarly, diverse cultures, religions, philosophies, and scientific investigations consider effects of vitality, conatus, élan vital, self-preservation, repair, redundancy, defense, nutrition, elixirs, or similar phenomena on mortality and survivorship^22,27,28,31,35,60,61,63,83,109-122^. The universality of tetraeffective causes of mortality and survivorship means that it is useful, practical, moral, and ethical to assume that every vitality is accompanied by an opposite frailty and vice versa, every damage is accompanied by an opposite repair and vice versa, every injury or disease is accompanied by an opposite remedy and vice versa, and so on; further implying that it is invalid, impractical, immoral, unethical, and not useful to assume that positive affects on mortality and negative effects on survivorship are unopposed; further implying that it is invalid, impractical, immoral, unethical, and not useful to assume that negative effects on mortality and positive effects on survivorship are unopposed. Thus, utilitarian, practical, moral, and ethical considerations provide additional rationales for the universality of tetraeffective causes of mortality and survivorship.

Previous scientific research does not provide mathematical specifications of tetraeffective causes, mortacauses, and vitacauses of mortality and survivorship. Moreover, previous scientific research does not provide evidence of mortacauses and vitacauses of tetraeffective causes of mortality and survivorship. Here I specify tetraeffective causes, mortacauses, and vitacauses, and I analyze humans’ and medflies’ mortality and survivorship in search of evidence of tetraeffective causes, mortacauses, and vitacauses.

## Methods

A previous investigation presents multivariable regression analyses of (1) 188,087 weighted cases with 79,164,608 events of death or survival of all individuals that were born in Sweden in decennial years 1760 – 1930 and died between 1760 and 2008, and (2) 50,716 weighted cases with 2,211,782 events of death or survival of caged Mediterranean fruit flies, *Ceratitis capitata*, commonly known as medflies^123^. These analyses employ AIC and BIC information criteria in tests of the following multivariable individual-level longitudinal limited powered polynomials binary random-effects regression model:

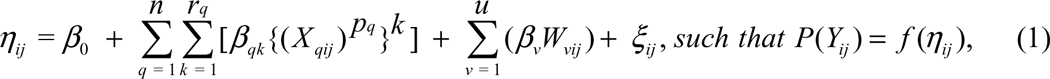

where *Y_ij_* denotes mortality *M_ij_* or survivorship *S_ij_* of an individual human or medfly *i* that continues to exist (i.e., *M_ij_* = 0 and *S_ij_* = 1) or ceases to exist (i.e., *M_ij_* = 1 and *S_ij_* = 0) at humans’ year *j* or medflies’ day *j*, *P(Y_ij_)* denotes the probability of mortality (i.e., *P(M_ij_))* or the probability of survivorship (i.e., *P(S_ij_)* of individual *i* at observation *j*, *f(η_ij_)* is a link function that denotes a transformation of *η_ij_* (e.g., a logit transformation *P(Y_ij_)* = *exp(η_ij_)/{1 + exp(η_ij_)}*), ***β*** denote regression coefficients *β*, ***X_q_*** denote ordinal or higher-level variables *X*, and ***W_v_*** denote categorical variables *W*. The specific ***X_q_*** variables in this investigation are: *X* = *A* denotes humans’ or medflies’ age, *X* = *L* denotes humans’ or medflies’ lifespan, *X* = *C* denotes humans’ or medflies’ contemporary aggregate size (i.e., this size refers to the number of individuals whose age, sex, and location in time or space are identical to those of the criterion individual), *X* = *Λ* denotes humans’ or medflies’ lifespan aggregate size (i.e., this size refers to the number of individuals whose lifespan, age, sex, and location in time or space are identical to those of the criterion individual), and *X* = *H* denotes humans’ historical time (i.e., a specific year). The specific ***W_v_*** variables in this investigation are: *W* = *F* denotes being female in reference to humans’ or medflies’ sex, and *W* = *Q* denotes medflies’ respective cages. Coefficients *q* denote sequential indicators of *n* distinct variables *X*, *p_q_* denotes a power coefficient of variable *X_q_*, *k* are sequential indicators of the *r_q_* polynomial length of variable *X_q_*. Coefficients *v* denote sequential indicators of *u* distinct variables *W*, and each *ξ_ij_* denotes a random-effects coefficient corresponding to individual *i* at observation *j*. The previous investigation provides further information on these data and regression analyses^123^.

A regression coefficient *β_1_* denotes here the first regression coefficient that applies to an ordinal or higher level variable *X* in Model 1, and a regression coefficient *β_2_* denotes here the second regression coefficient that applies to this variable *X* in Model 1, allowing for additional – i.e., third or more – regression coefficients for this variable *X*. Relationship *sign(β_1_)* = -*sign(β_2_)* for an ordinal or higher level variable *X*in Model 1 indicates here that variable *X* is a tetraeffective cause of mortality and survivorship. Relationships *β_1_* > 0 and *β_2_* < 0 at *Y* = *M* and relationships *β_1_* < 0 and *β_2_* > 0 at *Y* = *S* for a variable *X* in Model 1 indicate here that *β_1_* is an indicator of effects of an X-specific mortacause, further indicating that *β_2_* is an indicator of effects of an X-specific vitacause. Relationships *β_1_* < 0 and *β_2_* > 0 at *Y* = *M* and relationships *β_1_* > 0 and *β_2_* < 0 at *Y* = *S* for a variable *X* in Model 1 indicate here that *β_1_* is an indicator of effects of an *X*-specific vitacause, further indicating that *β_2_* is an indicator of effects of an X-specific mortacause. Each additional respective coefficient *β* for a variable *X*in Model 1 indicates here another respective *X*-specific mortacause or vitacause. Such indications apply to every variable *X* in Model 1. Respective *X*-specific multivariable regression coefficients *β_1_* and *β_2_* denote here the respective multivariable regression coefficients for the following variables *X* in Model 1: Humans’ and medflies’ age (*X* = *A*), lifespan (*X* = *L*), contemporary aggregate size (*X* = *C*), and lifespan aggregate size (*X* = *Λ*) as well as humans’ historical time (*X* = *H*).

## Results

Table 1 reveals *sign(β_1_)* = -*sign(β_2_)* in the multivariable regression analyses of effects age, lifespan, contemporary aggregate size, or lifespan aggregate size on humans’ and medflies’ mortality and survivorship, as well as in the multivariable regression analyses of effects of historical time on humans’ mortality and survivorship. Relationships *sign(β_1_)* = -*sign(β_2_)* in Table 1 thus respectively indicate here that humans’ and medflies’ age, lifespan, contemporary aggregate size, and lifespan aggregate size are respective tetraeffective causes of respective humans’ and medflies’ mortality and survivorship. Relationships *sign(β_1_)* = -*sign(β_2_)* in Table 1 also indicate that humans’ historical time is a tetraeffective cause of humans’ mortality and survivorship. Additionally, Table 1 reveals relationships *β_1_* < 0 and *β_2_* > 0 at *Y* = *M*, and Table 1 also reveals relationships *β*_1_ > 0 and *β_2_* < 0 at *Y* = *S*; further revealing here that respective indicate effects of respective *X*-specific vitacauses – and further revealing here that respective *β_2_* indicate effects of respective *X*-specific mortacauses – on respective humans’ and medflies’ mortality and survivorship, where *X* denote humans’ and medflies’ age (*X* = *C*) and lifespan (*X* = *L*) and where *X* denote medflies’ contemporary aggregate size (*X* = *C*) and lifespan aggregate size (*X* = *Λ*). Moreover, Table 1 reveals relationships *β_1_* > 0 and *β_2_* < 0 at *Y* = *M*, and Table 1 also reveals relationships *β_1_* < 0 and *β_2_* > 0 at *Y* = *S*; further revealing here that respective *β_1_* indicate effects of respective *X*-specific mortacauses – and further revealing that respective *β_2_* indicate effects of respective *X*-specific vitacauses – on humans’ mortality and survivorship, where *X* denote humans’ contemporary aggregate size (*X* = *C*) and lifespan aggregate size (*X* = *Λ*). Furthermore, Table 1 reveals relationships *β_1_* < 0, *β_2_* > 0, and *β_3_* < 0 at *Y* = *M*, and Table 1 also reveals relationships *β_1_* > 0 and *β_2_* < 0, and *β_3_* > 0 at *Y* = *S*; further revealing here that respective *β_1_* and *β_3_* indicate effects of respective humans’ historical-time-specific vitacauses – and respective *β_2_* indicates effects of humans’ historical-time-specific mortacause – on humans’ mortality and survivorship.

**Table 1.** Values of *β_1_* and *β_2_* in best-fitting Models 1. *Y* = *M* denotes mortality, *Y* = *S* denotes survivorship, *X* = *A* denotes age, *X* = *L* denotes lifespan, *X* = *C* denotes contemporary aggregate size, *X* = *Λ* denotes lifespan aggregate size, and *X* = *H* denotes historical time. Respective low and high *β_1_* and *β_2_* values respectively denote low and high values of these coefficients at respective 95% confidence intervals. *β_3_* = -7.97E-10, low, *β_3_* = -8.12E-10, high *β_3_* = -7.82E-10 at humans’ *X* = *H* and *Y* = *M*. *β_3_* = 7.97E-10, low *β_3_* = 7.82E-10, high *β_3_* = 8.12E-10 at humans’ *X* = *H*and *Y* = *S*. Adapted from a previous investigation*^123^*.

| entities | $Y$ | $X$ | $p$ | $\beta_1$ | $\beta_2$ | low $\beta_1$ | high $\beta_1$ | low $\beta_2$ | high $\beta_2$ |
| --- | --- | --- | --- | --- | --- | --- | --- | --- | --- |
| humans | M | A | 0.16 | -1074.55 | 546.12 | -1076.92 | -1072.19 | 544.92 | 547.32 |
| humans | S | A | 0.16 | 1074.55 | -546.12 | 1072.19 | 1076.92 | -547.32 | -544.92 |
| medflies | M | A | 0.13 | -2648.52 | 1295.76 | -2681.20 | -2615.85 | 1279.76 | 1311.76 |
| medflies | S | A | 0.16 | 1402.49 | -706.62 | 1373.72 | 1431.25 | -721.29 | -691.95 |
| humans | M | L | 0.88 | -17.12 | 0.10 | -17.16 | -17.08 | 0.10 | 0.10 |
| humans | S | L | 0.88 | 17.12 | -0.10 | 17.08 | 17.16 | -0.10 | -0.10 |
| medflies | M | L | 0.98 | -16.67 | 0.09 | -16.88 | -16.46 | 0.09 | 0.10 |
| medflies | S | L | 0.94 | 19.24 | -0.12 | 18.82 | 19.65 | -0.12 | -0.11 |
| humans | M | C | 0.75 | 0.00623 | -4.39E-07 | 0.00619 | 0.00627 | -4.46E-07 | -4.33E-07 |
| humans | S | C | 0.75 | -0.00623 | 4.39E-07 | -0.00627 | -0.00619 | 4.33E-07 | 4.46E-07 |
| medflies | M | C | 1.02 | -0.00632 | 6.85E-07 | -0.00652 | -0.00612 | 6.54E-07 | 7.15E-07 |
| medflies | S | C | 1.02 | 0.00407 | -4.03E-07 | 0.00388 | 0.00427 | -4.32E-07 | -3.74E-07 |
| humans | M | $\Lambda$ | 0.3 | 6.18689 | -0.34869 | 6.16888 | 6.20490 | -0.34969 | -0.34768 |
| humans | S | $\Lambda$ | 0.3 | -6.18689 | 0.348686 | -6.20490 | -6.16888 | 0.34768 | 0.34969 |
| medflies | M | $\Lambda$ | 0.95 | -0.09025 | 0.000263 | -0.09327 | -0.08723 | 0.00025 | 0.00027 |
| medflies | S | $\Lambda$ | 0.88 | 0.11285 | -0.00049 | 0.10723 | 0.11846 | -0.00052 | -0.00046 |
| humans | M | H | 1.41 | -7.84E-03 | 1.92E-06 | -7.91E-03 | -7.77E-03 | 1.86E-06 | 1.97E-06 |
| humans | S | H | 1.41 | 7.84E-03 | -1.92E-06 | 7.77E-03 | 7.91E-03 | -1.97E-06 | -1.86E-06 |

## Discussion

The foregoing considerations provide identifications, names, recognitions, elucidations, conceptions, specifications, explanations, and demonstrations of tetraeffective causes, mortacauses, and vitacauses of mortality and survivorship. These considerations of tetraeffective causes of mortality and survivorship usefully elucidate – and deepen the consideration of, and expand the scope of scientific research on – causes of mortality and survivorship. These considerations also provide a new paradigm of causes of mortality and survivorship, and enable and promote further scientific research and practical applications^124,125^. These considerations – and the methodology that is employed here and in a previous investigation^123^ – could thus prove to be particularly useful because scientific research on causality remains problematic and challenging^35,126-130^, scientific research on causes of mortality and survivorship remains particularly problematic and challenging^21,131-138^, and mortality and survivorship and their interrelationships are particularly prone to elicit errors and biases^139-144^.

Conceptions of deep or latent structures are found in diverse fields of science and scholarship^66,67,130,145-149^; conceptions of mortacauses and vitacauses of tetraeffective causes of mortality and survivorship have obvious affinities with conceptions of deep or latent structures, but much remains to be learned about these affinities. Additionally, as noted, conceptions of frailty, damage, disease, injury, waste, harm, poison, thanatos, destroyer of worlds, nuclear holocaust, global warming, poverty, injustice, and related phenomena are found in diverse cultures, religions, philosophies, and scientific investigations^27,28,60-63,93-108^; these conceptions have obvious affinities with conceptions of mortacauses, but much remains to be learned about these affinities. As also noted, conceptions of vitality, conatus, élan vital, self-preservation, repair, redundancy, defense, nutrition, elixirs, and related phenomena are found in diverse cultures, religions, philosophies, and scientific investigations^22,27,28,31,35,60,61,63,83,109-122^; these conceptions have obvious affinities with conceptions of vitacauses, but much remains to be learned about these affinities. Moreover, the extensive and longstanding considerations of oppositions in religion and philosophy^35,36,51-63^, quantum theory ^64,65^ structuralism^66-68^, biology ^69-71^, and art ^72-77^ emphasize that opposites differ with respect to specific variable or invariant characteristics (e.g., effectiveness, dominance, dynamism, intensity, potency, force, and other characteristics), further emphasizing that oppositions can be variable or invariant, and further emphasizing that oppositions are somehow resolved or come to some kind of *aufheben*^54,150,151^. These considerations – and the differences and similarities among effects of age, lifespan, contemporary aggregate size, lifespan aggregate size, and historical time on mortality and survivorship of humans and medflies in this investigation – show that much remains to be learned about the modulation, functions, evolutionary contexts, and characteristics of diverse kinds of tetraeffective causes, mortacauses, and vitacauses of mortality and survivorship of diverse kinds of entities in diverse times and places.

Searches for models and laws of mortality or survivorship are longstanding and inconclusive^22,27,31,42-46,80,100-102,115,116,120,152-172^. The considerations in this article provide a foundation for a hypothetical law of tetraeffective causes of mortality and survivorship; this hypothetical law states that every cause of mortality and survivorship is a tetraeffective cause that is composed of corresponding at least one cause-specific mortacause and at least one cause-specific vitacause. Specifications *sign(β_1_)* = -*sign(β_2_)* in multivariable regression models provide succinct, parsimonious, simple, and meaningful specifications of tetraeffective causes, mortacauses, and vitacauses of mortality and survivorship. Specifications *sign(β_1_)* = -*sign(β_2_)* and the hypothetical law of tetraeffective causes of mortality and survivorship contribute to the longstanding and inconclusive searches for models and laws of mortality and survivorship. The scope of these specifications and hypothetical law can be investigated in further research on diverse kinds of tetraeffective causes, mortacauses, and vitacauses of mortality and survivorship of diverse kinds of entities in diverse times and places^173,174^.

